# Modeling foot-and-mouth disease dissemination in Brazil and evaluating the effectiveness of control measures

**DOI:** 10.1101/2022.06.14.496159

**Authors:** Nicolas C. Cardenas, Francisco P. N. Lopes, Alencar Machado, Vinicius Maran, Celio Trois, Felipe Amadori Machado, Gustavo Machado

**Affiliations:** Department of Population Health and Pathobiology, College of Veterinary Medicine, North Carolina State University, Raleigh, NC, USA; Departamento de Defesa Agropecuária, Secretaria da Agricultura, Pecuária e Desenvolvimento Rural, Porto Alegre, Brazil; Laboratory of Ubiquitous, Mobile and Applied Computing (LUMAC), Polytechnic College of Federal University of Santa Maria, Santa Maria, Brazil; Department of Chemical Engineering, Rovira i Virgili University, Tarragona, Spain

**Keywords:** dynamical models, infectious disease control, epidemiology, transmission, targeted control

## Abstract

Foot-and-mouth disease (FMD) is known to infect multiple food-animal species and to spread among ungulate populations. We introduce a multiscale compartmental stochastic model that incorporates population dynamics, births, deaths, and species-specific transmission dynamics. The model integrates disease dynamics at both the between-farm and the within-farm levels. We developed four scenarios to evaluate the effectiveness of countermeasures, including movement standstill, vaccination, and depopulation. The base scenario involved vaccinating the animals on 20 farms and depopulating four infected farms. The three alternative control scenarios included increasing vaccination and depopulation capacities and no vaccination. Our findings indicate that bovines were the most frequently infected species, followed by swine and small ruminants. After ten days of initial spread, the number of infected farms ranged from 1 to 123, with most simulations (90.12%) predicting fewer than 50 infected farms. Most of the secondary spread occurred within a range of 25 km. Early response to initiating the control action reduces the number of days spent working on control actions while intensifying daily depopulation and vaccination capacity; this may be worth considering in decision-making processes for future control of FMD. Emergency vaccination proved to be efficient in reducing the magnitude and duration of outbreaks, whereas increasing depopulation without the use of vaccination also proved to be effective in eliminating the outbreaks.

## 1 Introduction

Foot-and-mouth disease (FMD) is an infectious disease of cloven-hoofed animals that affects multiple species, including bovines, swine, small ruminants, and wildlife (1). During the 2001 FMD epidemic in the U.K. and the Netherlands, more than 6.7 million animals were slaughtered, including healthy ones (preemptive culling) (2). In both outbreaks, multiple species were infected, including goats on mixed dairy goat/veal calf farms (2,3), and there were additional costs to other sectors, such as tourism, with a total expenditure of approximately 2.7 to 3.2 billion euros (4). The distribution of FMD outbreaks in South America has changed over the years; the most recent outbreaks occurred in 2017 and 2018 in Colombia (5), and no epidemics have been officially reported in Latin America since then, despite Venezuela’s absence of official international status for FMD (6). The official World Organization for Animal Health (WOAH) database for 2022 registered more than 57,000 outbreaks in 99 counties from 1996 to 2012; 68% of the FMD cases were associated with cattle, 22% with swine, and 21% with small ruminants and other species, whereas buffalo were not directly mentioned (7). Despite substantial evidence that all susceptible species can contribute to significant FMD epidemics, response plans frequently focus on controlling the spread of FMD among cattle populations. This approach often overlooks the role of other domestic species (8). For example, the large-scale 2001 UK outbreak was linked to swine slaughterhouses (9,10). Similarly, the most recent major outbreak in South America in 2001 affected 2,027 farms in Uruguay, where cattle and small ruminants were the predominant infected species (11).

Notably, FMD pathogenesis and transmission dynamics vary among species, given differences in the viral loads needed to cause infection and variability in the latency and duration of infection (12–15). For example, infected swine shed more viral particles than do cattle and sheep, historically resulting in widespread epidemics expected when swine are infected (14). Thus, it is pivotal to consider heterogeneity in transmission dynamics when modeling within- and/or between-farm FMD dissemination (8). Similarly, field observations and experimental trials have demonstrated that the spread of FMD between farms occurs primarily through direct contact between susceptible and infected animals (12,16) and via indirect contact with fomites and long-distance transport of aerosols, a process known as spatial transmission (16).

Mathematical models have been extensively utilized to investigate the propagation of FMD epidemics (17,18). Despite significant technical and computational advancements, simplifications of complex dynamics are often necessary due to computational costs or the lack of detailed population data, such as herd structure (e.g., number of individuals, number of animals born alive) and animal movements (17). A common simplification in modeling involves restricting the dynamics to a single species (4,19,20). Despite the varied applications and efforts in modeling FMD, outstanding questions persist regarding the measurement of epidemic trajectories and control strategies, particularly given the heterogeneous transmission dynamics among different susceptible species coexisting on the same premises (8,18).

Here, we develop a multihost, single-pathogen, multiscale model designed to capture the dynamics of FMD transmission patterns across different host species. The model i) simulates the spread of FMD within the state of Rio Grande do Sul in Brazil; ii) describes the geodesic distances between the initial outbreak and secondary cases; and iii) compares the effects of control action strategies, including emergency vaccination, depopulation, various restrictions on between-farm movement, and surveillance activities within control zones, taking into account the initial number of infected farms at the onset of control measures.

## 2 Materials and methods

This section presents the data used to create the dataset that was used to test and evaluate the model in outbreak simulations. The way in which the model was formulated is also described in this section, as are its outputs and the software used in its development.

### 2.1 Data sources

#### 2.1.1 Population data

A comprehensive dataset was compiled from the official records of 355,676 farms registered in Rio Grande do Sul, Brazil (21), hosted in the Agricultural Defense System (SDA)(22). The dataset included the number of animals per location individually for cattle, buffalo, swine, sheep, and goats. Following stringent criteria, 70,853 premises were excluded due to missing geographical coordinates, instances without animal stock, and the absence of incoming and/or outgoing movement during the study period spanning August 24, 2022 to August 24, 2023. Consequently, the final dataset included 284,823 farms with accurate and reliable information. To simplify the analysis, population and movement data from cattle and buffalo farms were merged into a single category denoted “bovines”. Similarly, sheep and goats were classified as “small ruminants”. Supplementary Material Figure S1 presents the farm-level population distribution, and Supplementary Material Figure S2 presents the geographical farm density distribution.

#### 2.1.2 Birth and death

Producers are required to disclose to the SDA at least once a year the total number of animals born alive on their farms and the number of deaths, including those due to natural causes. We collected data on births and deaths from the SDA; these data included 273,787 individual records associated with births and 268,790 records associated with deaths. The daily numbers of births and deaths, categorized by species, are depicted in Supplementary Material Figure S3.

#### 2.1.3 Movement data

From August 24, 2022 to August 24, 2023, 763,448 unique between-farm and farm-to- slaughterhouse movements were recorded; these were collected from the SDA centralized traceability database. Upon evaluation of the movement data, 106,481 records (13.9%) were removed for various reasons: a) lack of identification of origin or destination; b) zero animals moved; c) lack of specification of exact origin and destination premises; and d) movements to or from premises not registered in the population data or to premises outside the state of Rio Grande do Sul. Ultimately, 413,939 unique between-farm movements and 243,028 farm-to-slaughterhouse movements were analyzed. The daily farm-to-farm and farm-to-slaughterhouse movements, categorized by species, are depicted in Supplementary Material Figure S3.

### 2.2 Outbreak simulation

Over two hundred thousand farms are located in Rio Grande do Sul; from these, a sampling was made, and the sample farms were used as the initial infected premises. Our sample was multistage and stratified, with the number of farms and species categorized by municipality (23). The sample size was determined considering a prevalence of 50%, with a 95% significance level and a 1.1% margin of error. The sampling yielded a total of 10,294 farms (Supplementary Material Figure S4). Our model simulation was performed in two steps. First, FMD was seeded randomly into sample farms via one infected animal between August 24, 2022 and August 24, 2022. For farms with multiple species, for example, those with bovines, swine, and small ruminants, FMD was seeded into bovines; for farms with swine and small ruminants, FMD was seeded into swine; and for farms with cattle and small ruminants, FMD was seeded via one infected bovine. We assumed that all the animals were susceptible to FMD, as the annual vaccination campaign in Rio Grande do Sul had been suspended since May 2021(24).

### 2.3 Model formulation

We developed a multihost, single-pathogen, coupled multiscale model to simulate FMD epidemic trajectories (25) and subsequently applied countermeasures. The model led to the development of an R and Python package entitled “MHASpread: A multihost animal spread stochastic multilevel model” (version 0.3.0); additional details can be found at https://github.com/machado-lab/MHASPREAD-model. MHASpread allows the explicit specification of species-specific transmission probabilities and the latent and infectious periods of a disease that infects multiple species. The within-farm level includes births and deaths for each species. The between-farm level includes the entry and exit of animals due to between-farm movements and movements to slaughterhouses (Figure 1).

**Figure 1.**
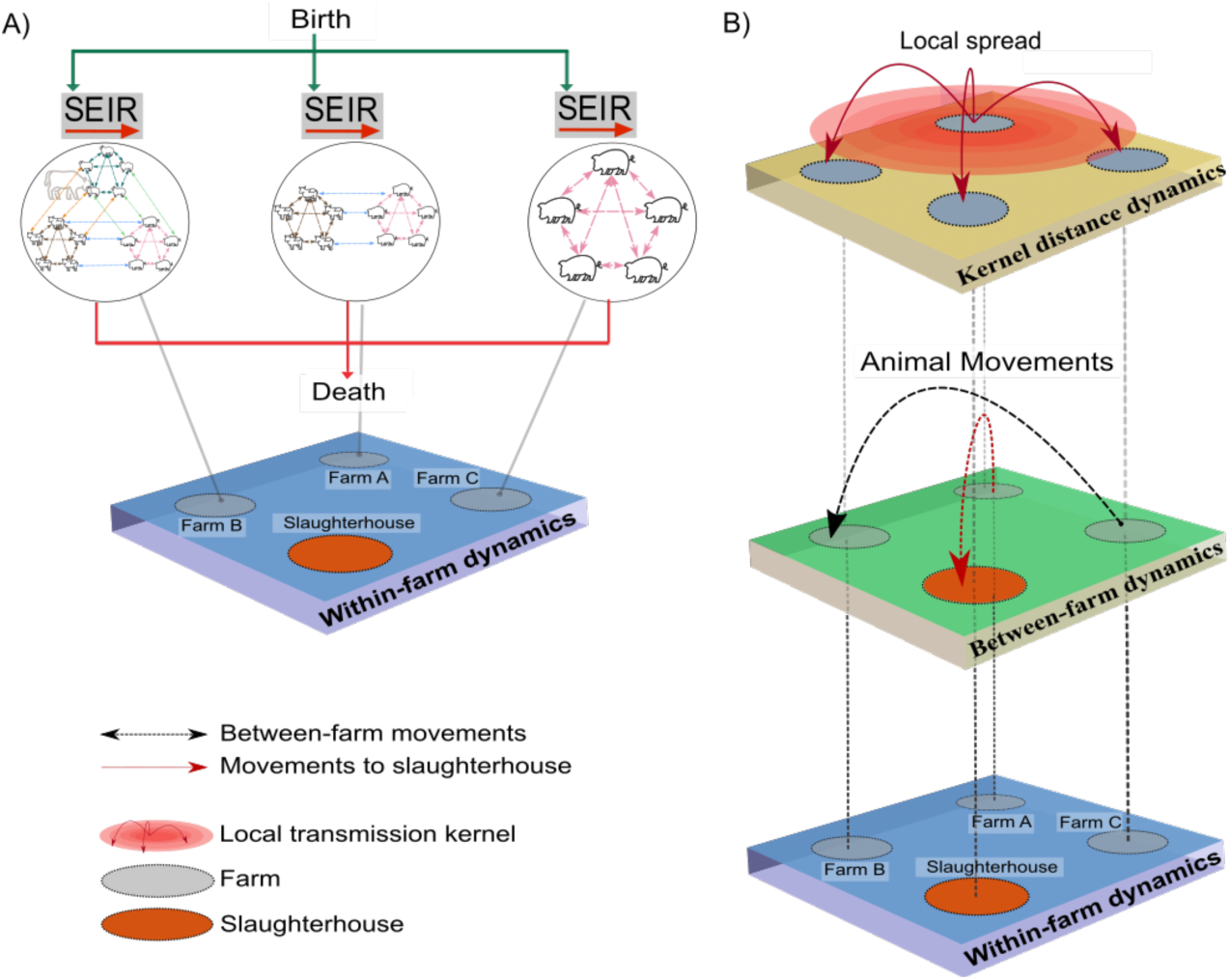
Schematic of state transitions in within-farm and between-farm dynamics. A) Within- farm dynamics: The green arrows indicate the introduction of animals (births) into the susceptible (S) compartment at farm *i* at time *t*. Each circle indicates a farm containing single or multiple species with specific host-to-host transition parameters (σ, γ); the dashed lines represent interactions within and between host species. The red arrows represent the removal of animals (deaths) regardless of infection status. B) Between-farm dynamics: The layer represents the number of animals moved (batches; n) from the farm of origin (*i*) to a destination farm (*j*) at time t (indicated by the black dashed arrows). Animals that moved to the slaughterhouse were removed from the simulation regardless of their infection status and are indicated by red dashed arrows. The kernel distance dynamics represent the spatial transmission distances.

#### 2.3.1 Within-farm dynamics

For within-farm dynamics, we assume that populations are homogeneously distributed. Species were homogeneously mixed in farms with at least two species, meaning that the probability of contact among species was homogeneous regardless of whether species were segregated by barns (e.g., commercial swine farms are housed in barns with limited changes in direct contact with cattle). The within-farm dynamics consist of mutually exclusive health states (i.e., an individual can only be in one state per discrete time step) for animals of each species (bovines, swine, and small ruminants). These health states (hereafter, “compartments”) include susceptible (*S*), exposed (*E*), infectious, (*I*), and recovered (*R*) and are defined as follows:

i. Susceptible: animals that are not infected and are susceptible to infection.
ii. Exposed: animals that have been exposed but are not yet infected.
iii. Infectious: infected animals that can successfully transmit the infection.
iv. Recovered: animals that have recovered and are no longer susceptible.

Our model considers birth and death, and these parameters are used to update the population of each farm. The total population is calculated as *N* = *S* + *E* + *I* + *R*. The number of individuals within each compartment transitions from *Sβ* → *E*, 1/*σ* → *I*, 1/*γ* → *R* according to the following equations:

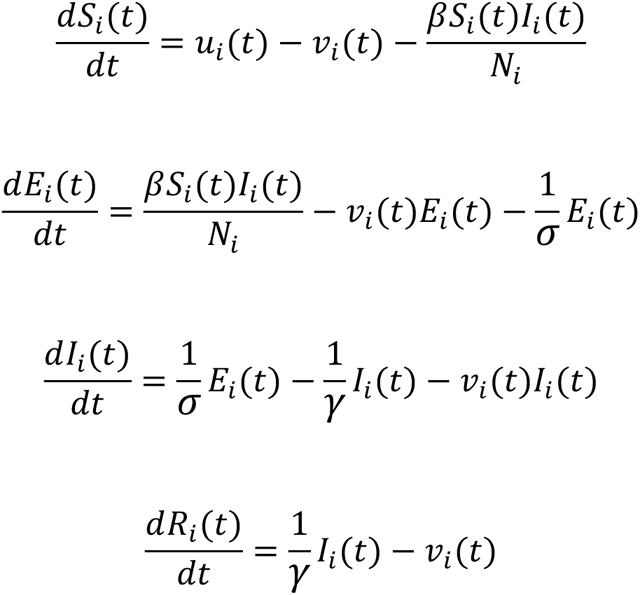

Transmission depends on the presence of infected and susceptible host species, as reflected by the species-specific FMD transmission coefficient *β* (Table 1.) Births are represented by the number of animals born alive *v_i_*(*t*) that enter the *S* compartment on the farm *i* at time *t* according to the day-to- day records; similarly, *v_i_*(*t*) represents the exit of the animals from any compartment due to death at time *t*. The transition from *E* to *I* is driven by 1/*σ*, and the transition from *I* to *R* is driven by 1/*γ*; these values are drawn from the distribution generated for each specific species according to the literature (Table 2). The within-farms dynamics are depicted in Figure 1.

**Table 1.** Distribution of host-to-host transmission coefficients (β) per animal^-1^ day.

| Infected species | Susceptible taxon | Transmission coefficient ($\beta$), shape and distribution | Reference |
| --- | --- | --- | --- |
| Bovines | Bovines | PERT (0.18, 0.24, 0.56) | Calculated based on the 2000-2001 FMD outbreaks in the state of Rio Grande do Sul (24). |
| Bovines | Swine | PERT (0.18, 0.24, 0.56) | Assumed. |
| Bovines | Small ruminants | PERT (0.18, 0.24, 0.56) | Assumed. |
| Swine | Bovines | PERT (3.7, 6.14, 10.06) | Assumed (26). |
| Swine | Swine | PERT (3.7, 6.14, 10.06) | (26). |
| Swine | Small ruminants | PERT (3.7, 6.14, 10.06) | Assumed (26). |
| Small ruminants | Bovines | PERT (0.044, 0.105, 0.253) | Assumed (27). |
| Small ruminants | Swine | PERT (0.006, 0.024, 0.09) | (28). |
| Small ruminants | Small ruminants | PERT (0.044, 0.105, 0.253) | (27). |

**Table 2.**
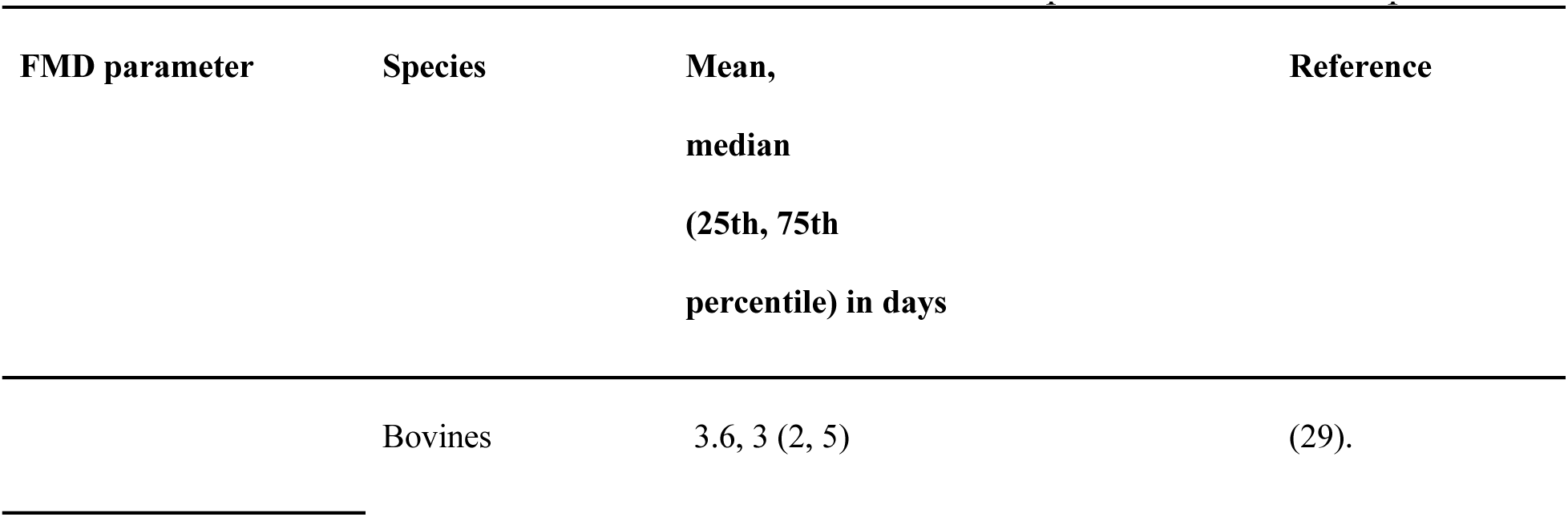
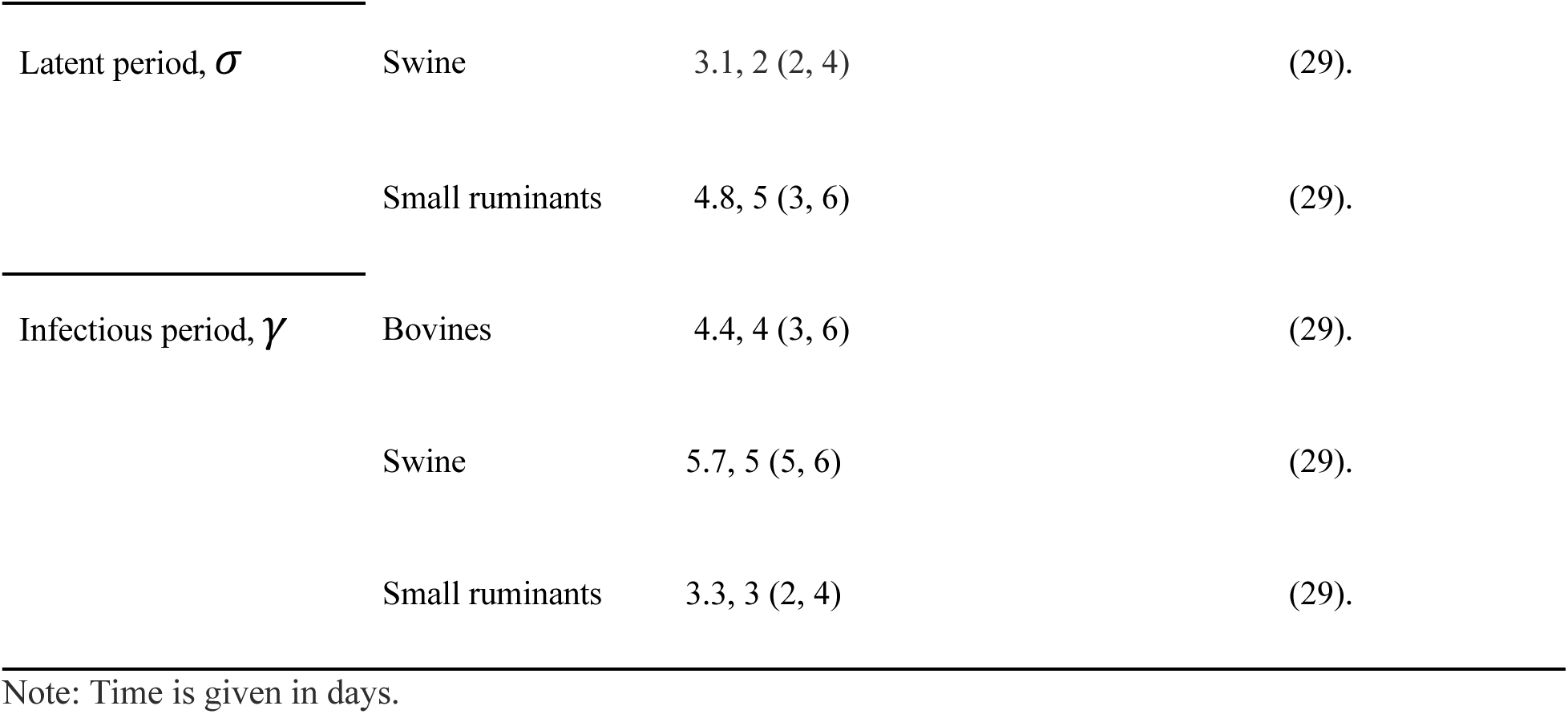
Within-farm distributions of latent and infectious FMD parameters for each species.

| FMD parameter | Species | Mean, median (25th, 75th percentile) in days | Reference |
| --- | --- | --- | --- |
|  | Bovines | 3.6, 3 (2, 5) | (29). |
| Latent period, $\sigma$ | Swine | 3.1, 2 (2, 4) | (29). |
|  | Small ruminants | 4.8, 5 (3, 6) | (29). |
| Infectious period, $\gamma$ | Bovines | 4.4, 4 (3, 6) | (29). |
|  | Swine | 5.7, 5 (5, 6) | (29). |
|  | Small ruminants | 3.3, 3 (2, 4) | (29). |

Note: Time is given in days.

#### 2.3.2 Kernel transmission dynamics

Spatial transmission encompasses a range of mechanisms that include airborne transmission, animal contact over fence lines, and sharing of equipment between farms (30,31). Local spread was fitted via a spatial transmission kernel in which the likelihood of transmission decreased as a function of the between-farm distance. The probability *PE* at time *t* describes the likelihood that the animals on a farm become exposed and is calculated as follows:

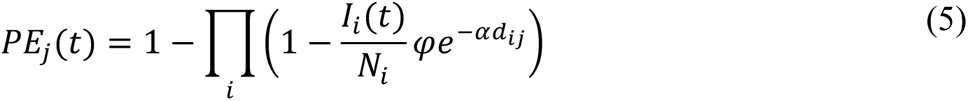

where *j* represents the uninfected population and where *ad_ij_* represents the distance between farm *j* and infected farm *i*, within a maximum distance of 40 km. Given the extensive literature on distance-based FMD dissemination and a previous comprehensive mathematical simulation study (32), distances above 40 km were not considered. Here, 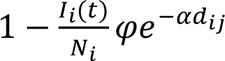 represents the probability of transmission between farms *i* and *j* scaled by the prevalence of infection on farm 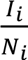, given the sdistance between the farms in kilometers (Figure 2). The parameters *φ* and *α* control the shape of the transmission kernel, *φ* = 0.044, which is the probability of transmission when *d_ij_* = 0, and *α* = 0.6 controls the steepness with which the probability decreases with distance (30,31). The relationship of exposure probability to distance is depicted in Figure 2.

**Figure 2.**
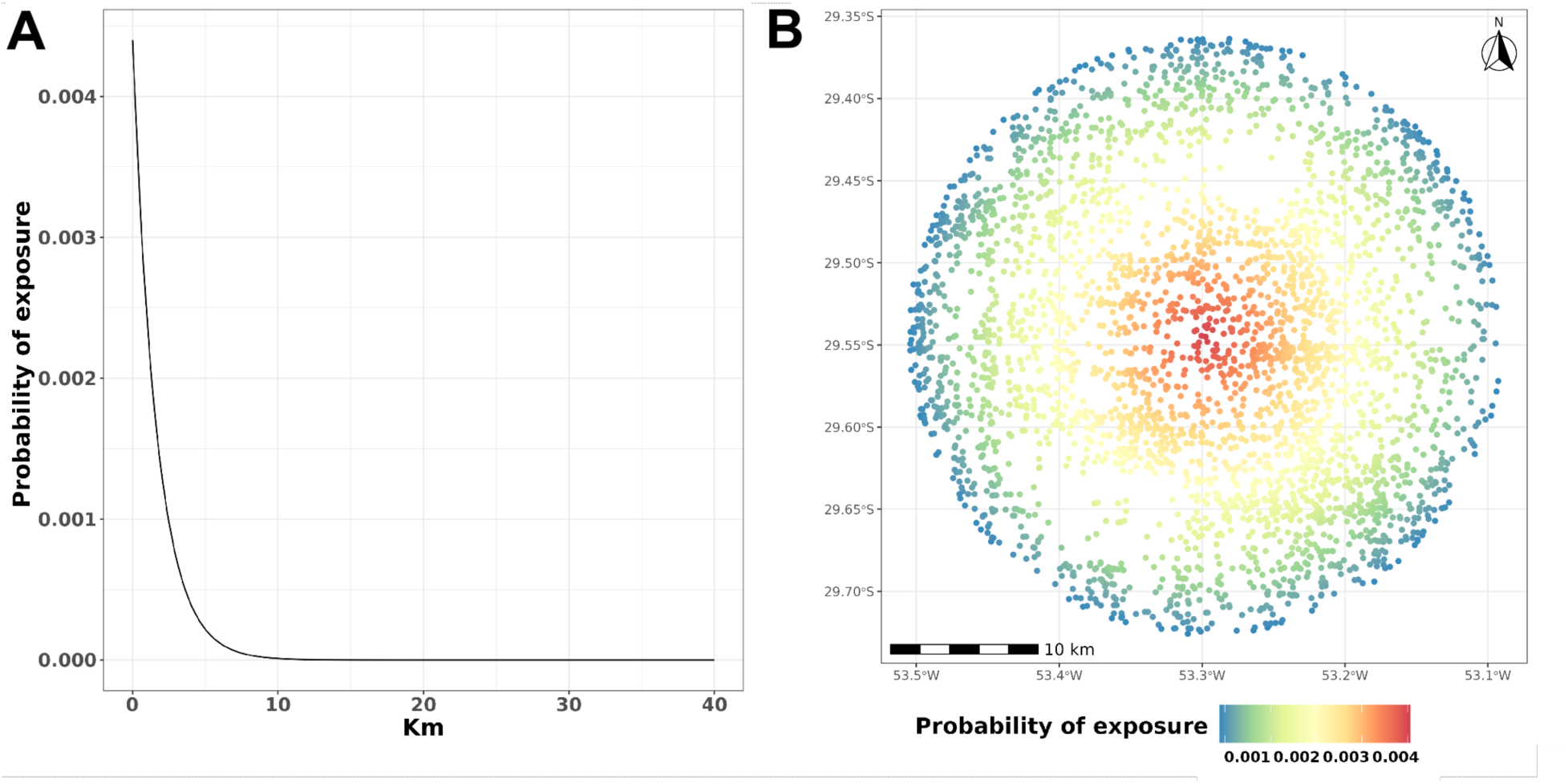
A) The y-axis represents the probability of exposure *PE*, and the x-axis represents the distance in km. B) Representation of farm locations. The color of each dot represents the probability of *PE* exposure; in this example, the infected farm is located in the center of the radius.

### 2.4 FMD spread and control actions

We first simulate an initial silent spread over ten days. This procedure yielded a wide range of initial outbreak scenarios, depicted in Figure 2, before the implementation of control actions. Next, we outline four different control action scenarios that were described by the Brazilian Ministry of Agriculture and Livestock (33):

*The baseline control scenario, which is referred to hereafter as “base”*, considers the following measures: i) depopulation of infected farms; ii) emergency vaccination of the animals on all farms in the infected and buffer zones; iii) restriction of animal movement for 30 days; and iv) establishment of three distinct control zones around infected farms at the following distances: 3-km infected zone, 7-km buffer zone, and 15-km surveillance zone (Supplementary Material Figure S5).

Depopulation: Involves the removal of all animals from infected farms located within the infected zone(s). For this study, the daily depopulation capacity was set to four farms, a value that aligns with the maximum depopulation capacity observed in Rio Grande do Sul (Dr. Fernando Groff, personal communication). Farms with relatively high animal populations were prioritized for depopulation. Once depopulated, these farms are no longer considered in the simulation. If the daily depopulation capacity was insufficient to cover all identified infected farms within one day, those farms were scheduled for depopulation on the following day or as soon as possible, respecting the maximum capacity constraints.

Emergency vaccination: The bovine farms within the infected and buffer zones are vaccinated. The daily maximum capacity is ten farms in the infected zone(s) and ten farms in the buffer zone(s) (Dr. Fernando Groff, personal communication). We simulated the delay in starting vaccination by setting vaccination to begin seven days post-FMD detection. Farms not vaccinated within a day due to the limited vaccination capacity were vaccinated on the subsequent day(s). Additionally, vaccine effectiveness was 90% in 15 days (34,35). For details about the implementation of emergency vaccination, Supplementary Material Control actions.

Traceability: We utilized contact tracing to identify farms that had direct contact with infected farms within the past 30 days. These farms underwent surveillance, including clinical examination of the animals and disease detection. Farms with animals that tested positive during traceback were categorized as infected farms.

Movement standstill: A 30-day restriction on animal movement across all three control zones was implemented; this restriction prohibited any incoming or outgoing movement of animals. The control zones were lifted, and the restriction on movement was maintained until depopulation was complete.

We assumed that we identified 10% of the infected farms when we began controlling them (10 days after introducing the index case). For example, if 100 farms were infected initially, 10 of those farms were found. When the number of detected farms was less than one, we rounded up to one detected farm. Furthermore, the rate of detection was influenced by two primary factors: the total number of farms within the control zones and the number of infected farms. For example, when fewer farms are under surveillance but the number of infected farms is higher, the likelihood of detection increases (Supplementary Material Figure S6). Infected farms located outside the control zones were also considered to be among the farms subject to detection.

#### 2.4.1 Alternative control scenarios

*Base x 2*: In this scenario, the daily number of vaccinations was increased to 40 farms, and depopulation was increased to eight farms. *Base x 3:* This scenario differed in that it assumed 60 farms vaccinated daily and 12 farms depopulated. *Depopulation*. This scenario differed from the baseline control scenario in that the number of infected farms depopulated was increased to 12 and vaccination was not used.

### 2.5 Model outputs

Our simulations tracked the number of animals in each health compartment and the number of infected farms at each time step. The epidemic trajectories were used to calculate the geodetic distances in km between the seeded index infections and the subsequent infections. In addition, we determined the probability of distance-dependent transmission by calculating the cumulative empirical spatial distribution. We utilized a generalized additive (GAM) model to plot the relationship between the number of infected farms and the number of days across different scenarios, as well as one-way analysis of variance (ANOVA) with a Tukey post hoc test to compare the scenarios. This enabled us to explore potential nonlinear relationships between the variables, effectively capturing complex patterns that might exist in the data. In addition, a mixed-effects regression model was fitted to describe the relationship between days working on control action and the initial number of infected farms controlled by each scenario.

### 2.6 Sensitivity analysis

We used a combination of Latin hypercube sampling (LHS), developed by (36), and the partial rank correlation coefficient (PRCC) technique to perform a local sensitivity analysis. LHS is a stratified Monte Carlo sampling method without replacement that provides unbiased estimates of modeling output measures subject to combinations of varying parameters. The PRCC approach can be used to classify how output measures are influenced by changes in a specific parameter value while linearly accounting for the effects of other parameters (37). As input model parameters, we selected the following categories and interspecies interactions: β bovine to bovine, β bovine to swine, β bovine to small ruminants, σ bovine, γ bovine, β swine to swine, β swine to bovine, β swine to small ruminants, σ swine, γ swine, β small ruminants to small ruminants, β small ruminants to bovine, β small ruminants to swine, σ small ruminants, and γ small ruminants. In total, 15 parameters were used to classify the monotonic relationship of infection status to our input variables to classify model sensitivity. The inputs include one farm at which the initial conditions were varied across 10,000 simulations over the LHS space. A positive PRCC indicates a positive relationship with the number of infected animals, whereas a negative PRCC indicates an inverse relationship with the number of infected animals; however, the magnitude of the PRCC does not necessarily indicate the importance of a parameter (38).

### 2.7 Software

The MHASpread model was developed, and graphics, tables, and maps were created using R v. 4.1.1 (39) and Python v. 3.8.12. Sampler (40), tidyverse (41), sf (42), brazilmaps (43), doParallel (44), and lubridate (45) were the packages used with R v. 4.1.1, and Numpy (46), Pandas (47), and SciPy (48) were the packages used with Python v. 3.8.12. The model is available in both R and Python versions.

## 3 Results

### 3.1 Initial spread and detection

Initially, we explored the variation in initial infection trends within the first ten days (Figure 3). The median number of infected farms was 52.5 (IQR: 26.75 to 78.25, maximum 123). The majority of the infected farms were swine farms (median 43.5, IQR: 22.25 to 64.75, maximum 105); the median number of infected bovine farms was 43 (IQR: 22 to 64, maximum 85), and the median number of infected small ruminant farms was 20.5 (IQR: 10.75 to 30.25, maximum 42). The full distribution of the epidemic trajectories is depicted in Supplementary Figure S7.

**Figure 3.**
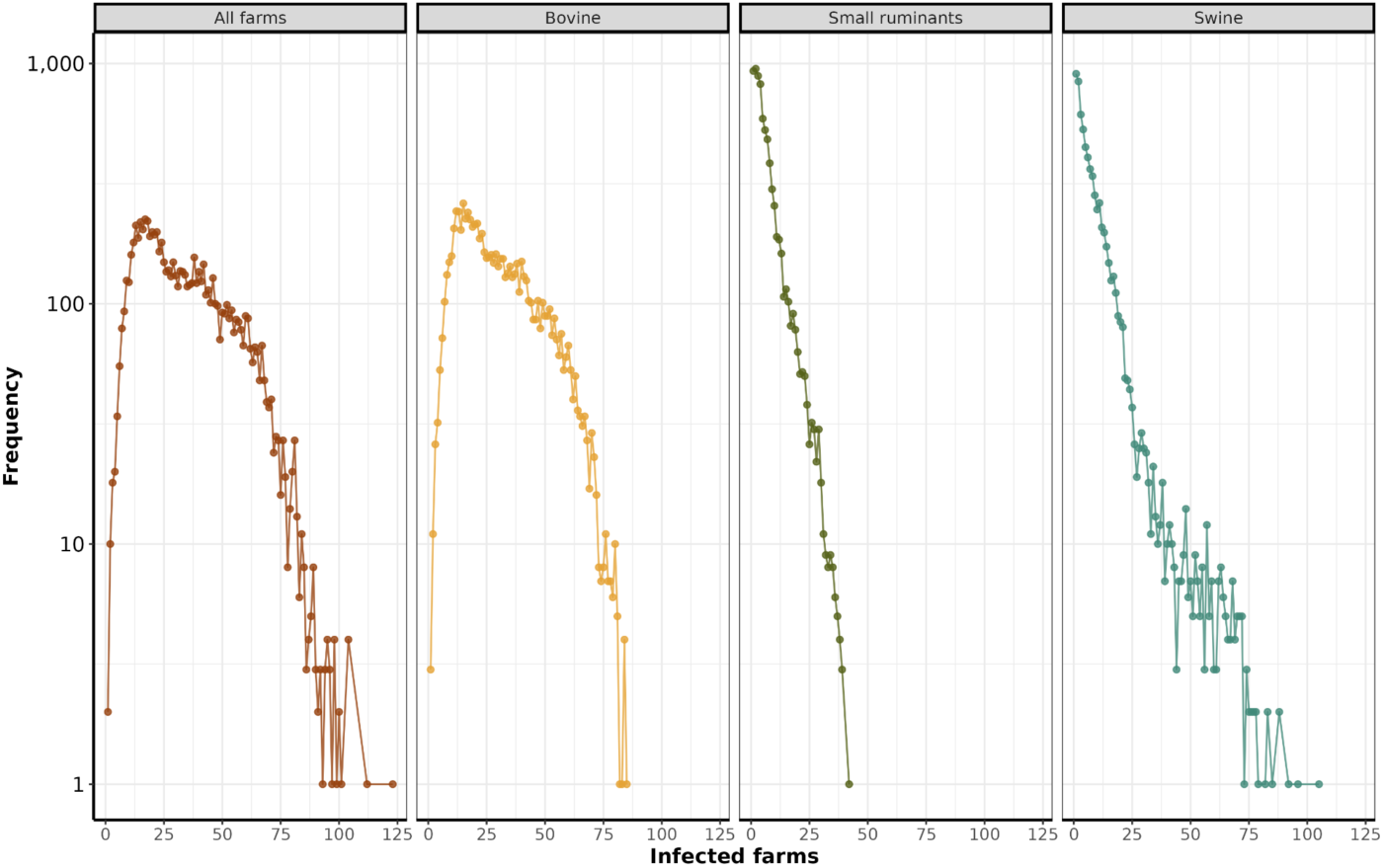
Distribution of the initial infected farms. The y-axis depicts the logarithm (base 10) of the frequency of infected farms across all simulations. The x-axis represents the number of infected farms.

### 3.2 Distances from the initial outbreak

Among the 284,396 unique simulated FMD trajectories, the median distance from the seeded infection site to the secondarily infected farm within the first ten days was 4.78 km (IQR: 2.64 km to 7.98 km, maximum 6.88 km) (Figure 4 A). Furthermore, we observed a linear increase with time in the distance over which the FMD disseminated (Figure 4 B).

**Figure 4.**
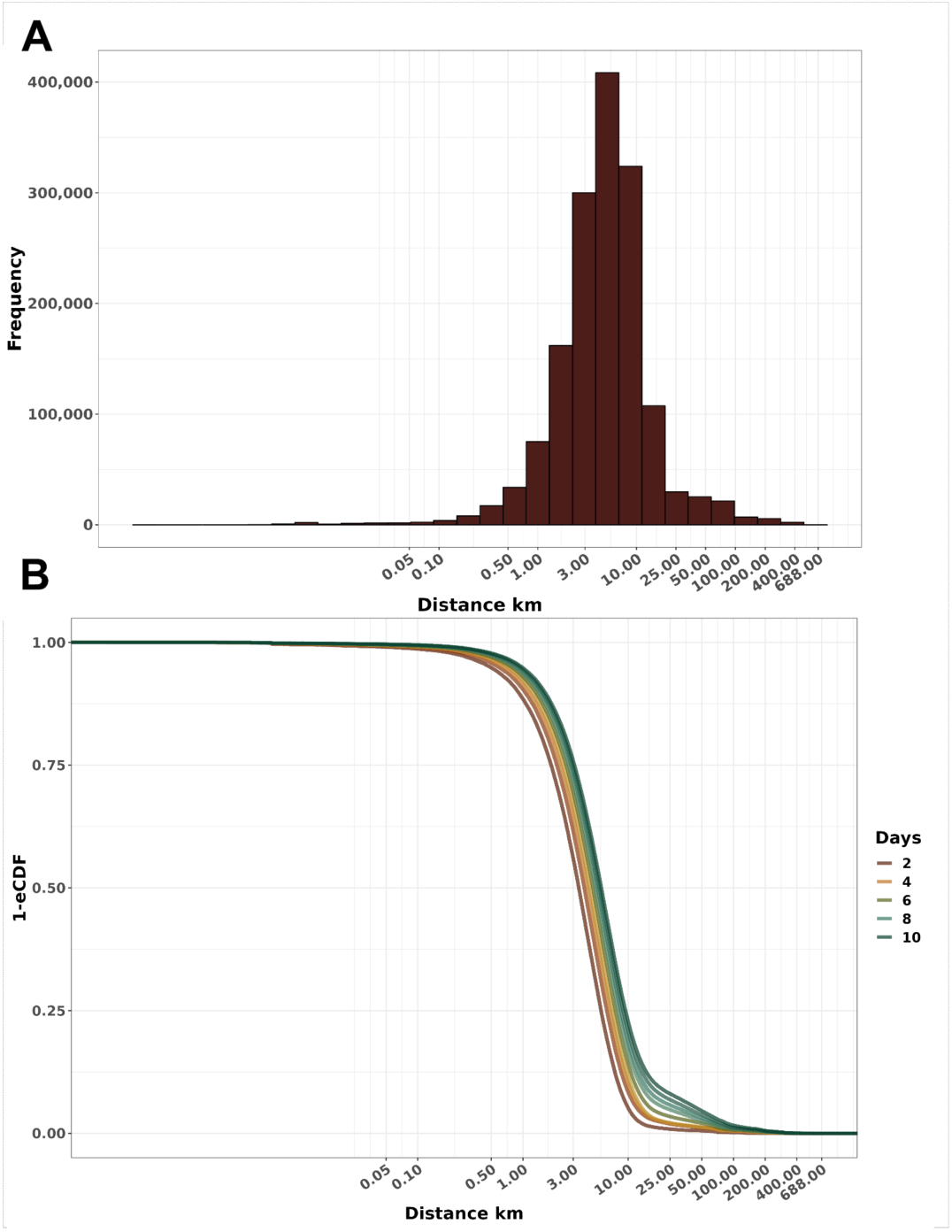
Distribution of the cumulative distance of dissemination. A) Frequency histogram of secondary infections at varying distances from the seeded infection. B) Empirical cumulative distribution function (1-ECDF) of the probability of infection according to the distance from the initial outbreak after disease introduction. Both x-axes are presented on a log10 scale.

### 3.3 Effectiveness of control measures

All the control scenarios were effective in eliminating all outbreaks within 120 days of the start of the control measures. However, their effectiveness was significantly different (ANOVA, p-value <0.05), except when we compared *depopulation* with the *base x 3* scenario. In general, the most effective alternative scenarios were *base x 3* and *depopulation*. The most notable differences in the average of infected farms by scenario were between the *base* and *depopulation* scenarios, with mean differences of 2.36 (95% CI: 2.22 to 2.49), followed by *base x 3* and *base* with 2.26 (95% CI: 2.13 to 2.39) and *base x 2* and *base* with 1.35 (95% CI: 1.22 to 1.49) (Supplementary Material Figure S8). In addition, we used a generalized additive model (GAM) to visualize the course of the simulated epidemics over time. Notably, scenario *base x 3* consistently exhibited a lower prevalence over time than the *depopulation*, *base x 2*, and *base* scenarios (Figure 5).

**Figure 5.**
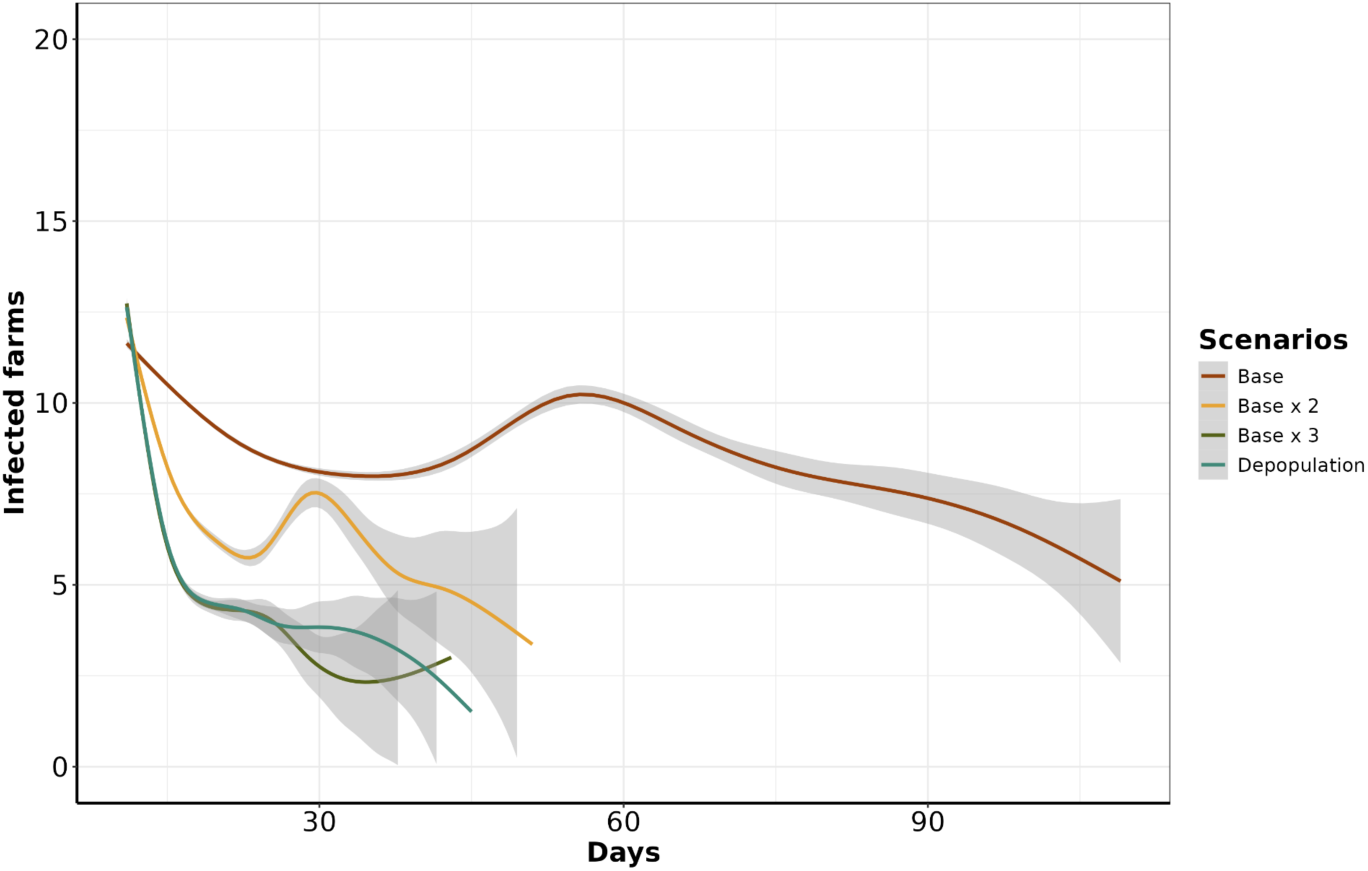
Estimated number of infected farms on days 11 through 120. The y-axis represents the number of infected farms, and the x-axis represents the days from start of simulation until the day with infected farms. The differently colored lines correspond to the individual scenarios.

#### 3.3.1 Duration of control actions

The median number of days over which the control actions were implemented for the *base* scenario was 22 (IQR: 17 to 29, maximum 109). While *base x 2* had a median of 16 days (IQR: 14 to 19, maximum 51), *base x 3* had a median of 15 days (IQR: 13 to 17, maximum 43), and *depopulation* had a median of 14 days (IQR: 13 to 17, maximum 45) (Figure 6). In addition, we measured the similarities and disparities between the mean number of days during which control actions were implemented, meaning the time period during which at least one outbreak response action was still ongoing. The comparisons between *depopulation* and *base*, *base x 3* and *base*, and *base x 2* and *base* revealed substantial disparities in the group means: -9.77 (95% CI: -10.04 to -9.49), -9.83 (95% CI: - 10.10 to -9.56), and -8.09 (95% CI: -8.36 to -7.81), respectively. A complete statistical analysis of these differences is presented in Supplementary Material Figure S9. Our findings indicate that there is a positive relationship between the time spent in control action and the number of farms infected at the beginning of the control actions. We found a linear relationship that indicates that, on average, for each additional farm that is infected at the beginning of the control actions, the number of days spent working on control actions increases by approximately 1.59 days (GLM, p-value < 0.05).

**Figure 6.**
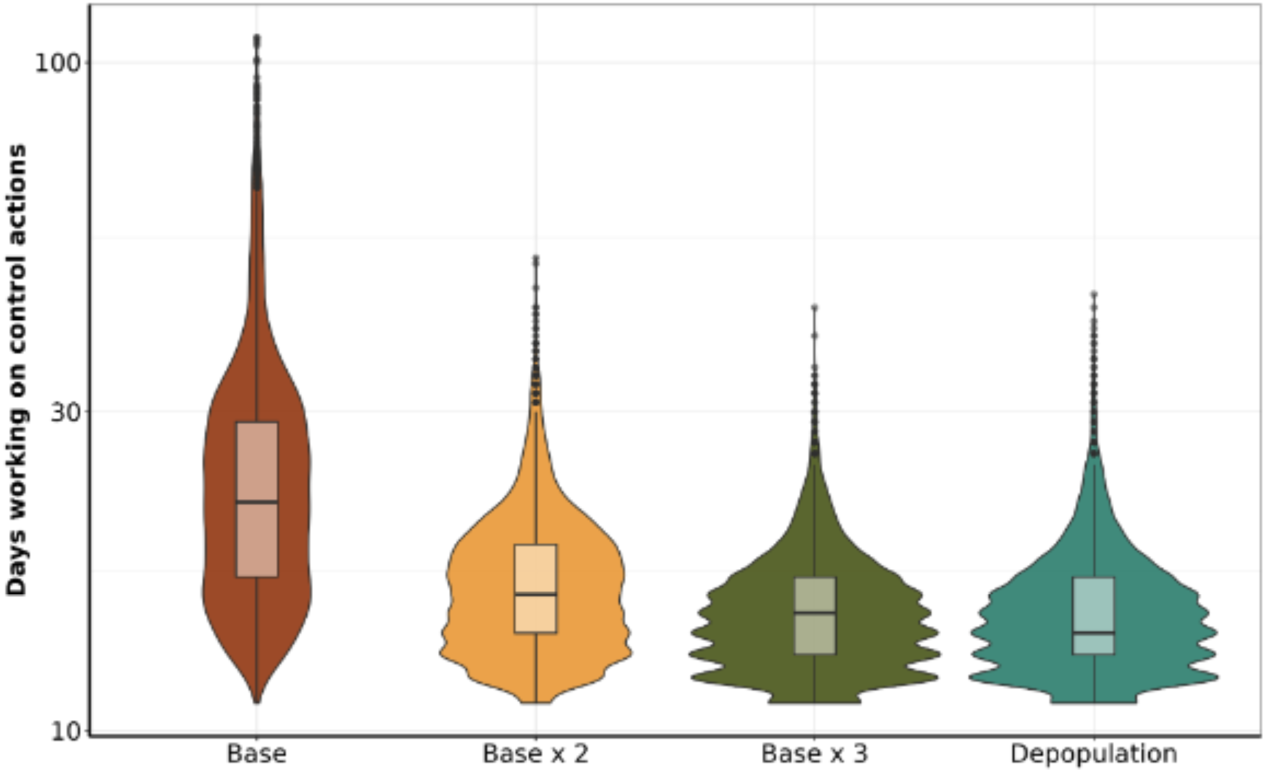
Comparison of the number of days spent working on control actions. The y-axis shows the total number of days dedicated to each control action, and the x-axis presents the scenarios.

#### 3.3.2 Vaccination

In the *base* scenario, the daily median number of vaccinated animals was 1,928 (IQR: 1,562 to 3,567, maximum: 20,740). In the *base x 2* scenario, the median increases to 3,959.32 (IQR: 3,067.75 to 5,865.56, maximum: 25,877). Similarly, in the *base x 3* scenario, the median increases to 5,947 (IQR: 4,157 to 8,384, maximum: 25,006). In the initial 30 days, there was a significant increase in the number of vaccinated animals, after which the amount of vaccine continued to increase with decreasing step demand (Figure 7).

**Figure 7.**
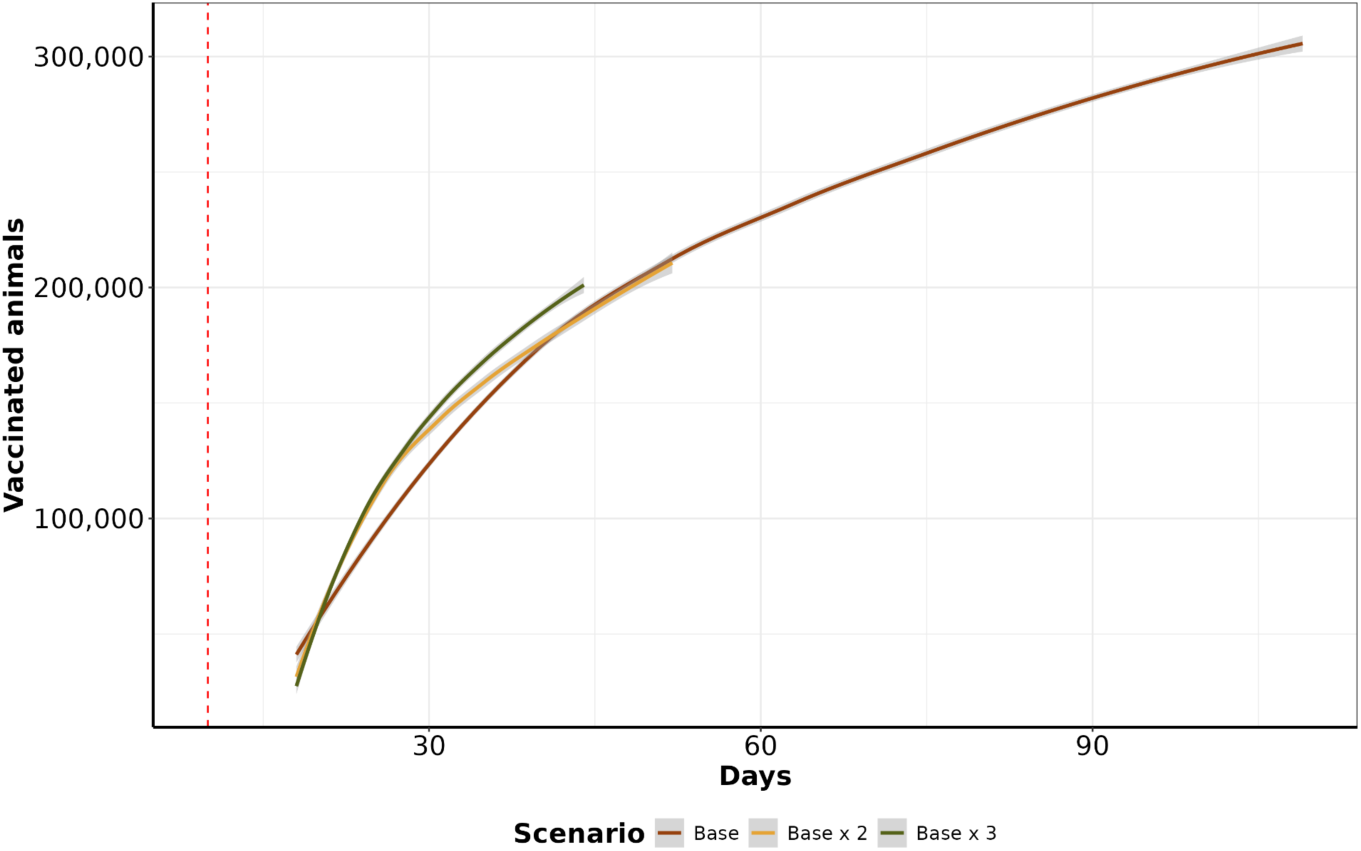
Vaccination curve by scenario. The y-axis represents the cumulative average number of vaccinated animals per day on a log10 scale. The x-axis shows the day on which the animals were vaccination. The red dashed line represents the curve obtained when control actions were initiated after 15 days of initial control actions.

#### 3.3.3 Depopulation

We analyzed the daily average number of depopulated animals over time. The scenarios *base x 2* and *depopulation* had the highest cumulative means of 3,071 (IQR: 1,767 to 3,768, maximum: 4,120) and 2,159 (IQR: 1,314 to 3,000, maximum: 3,830), respectively. The *base* and *base x 3* scenarios followed closely, with means of 2,139 (IQR: 1,798 to 2,541, maximum: 3,039) and 1,151.93 (IQR: 500 to 2,799, maximum: 4,398), respectively. The *depopulation* scenario consistently resulted in the highest count of affected animals, followed by the *base x 3*, *base x 2*, and *base* scenarios, particularly for bovines and small ruminants (Figure 8). However, in the case of swine, the *base* and *base x 2* scenarios presented higher numbers of depopulated animals than did the other species.

**Figure 8.**
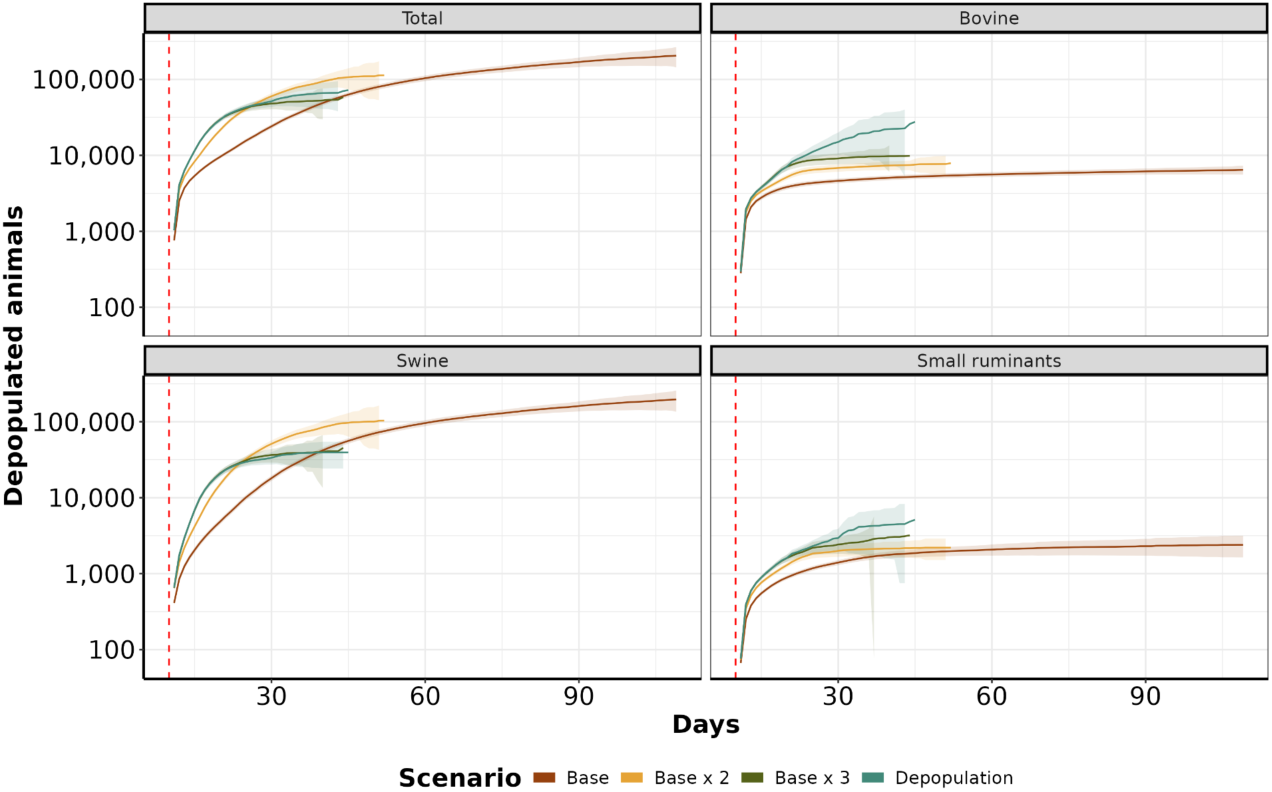
Depopulation curves by scenario. The y-axis shows the daily cumulative average number of depopulated animals. The x-axis represents the days since the start of control actions until the last depopulated farm. The red dashed line represents the day in which control actions started.

### 3.4 Sensitivity analysis

We evaluated the sensitivity of 15 model parameters and found them to have weights ranging from - 0.86 to 0.72; this sensitivity indicates a limited influence of the model parameters on the number of simulated secondary infections. Specifically, the length of the latent period, *σ*, had a negative impact on the number of secondary infections, and the length of the infectious period, *γ*, had a positive influence on the number of secondary infections; the results for both *σ* and *γ* were significant (p value <0.05) for the overall simulated species. The complete results of the sensitivity analysis are presented in Supplementary Material Figure S10.

## 4 Discussion

This study aimed to develop a multiscale, multihost stochastic model of FMD that explicitly incorporates species-specific transmission interactions. Our model was used to simulate the spread of FMD among cattle, buffalo, swine, and small ruminants in Rio Grande do Sul, Brazil, and to examine the effectiveness of countermeasures. Bovine farms were the most infected, followed by farms with swine and small ruminants, mostly because of the greater number of cattle farms and the connectivity of the swine contact network. Most secondary infections spread within a distance of 25 km, showing that proximity plays an important role in the dynamics of disease transmission. Our simulations demonstrated that tripling the number of daily depopulations and vaccinations eliminated epidemic trajectories within 15 days: this required the vaccination of 5,947 animals and the depopulation of 2,139 animals.

The number of farms infected within ten days of the introduction of the disease ranged from 1 to 123, with the majority of simulations (90.12%) resulting in fewer than 50 infected farms. Our study also revealed that when FMD infects swine, the sizes of the resulting epidemics are significant (Figure 3). This risk is particularly relevant to areas that harbor dense swine populations associated with commercial swine production; farms engaged in this type of production are typically vertically integrated, meaning that they move a significant number of swine, facilitating long-distance spread (49,50). Our study demonstrated that the number of farms initiating control actions has a linear effect on the duration of these actions, regardless of the implemented scenario. Specifically, each additional infected farm extended the duration of control actions by an average of 1.6 days. This indicates that enhancing the sensitivity of animal disease detection is crucial for optimizing the effectiveness of control strategies (51). Based on this finding, we argue that improving the timing of detection and optimizing the response to and the management of outbreaks are pivotal in ensuring effective control. Compared with the other proposed scenarios, scenario *base x 3* demonstrated the best performance, requiring a median of 15 days of control actions to eliminate the outbreak (Figure 4). In comparison, the *base* scenario required a median duration of 22 days, and the *base x 2* scenario required a median duration of 16 days. An average of 5,947 vaccine doses per day were administered in the *base x 3* scenario, compared with 1,928 per day in the *base* scenario and 3,959 per day in the *base x 2* scenario. Finally, the depopulation of 12 farms daily was successful in mitigating outbreaks; however, this scenario poses a significant challenge for official services.

Emergency vaccination presents an alternative to preemptive culling but may also limit the ability of surveillance systems to detect infected farms, as it may mask clinical signs of disease (52,53). When the median number of vaccinated animals at the end of the control action scenarios was examined, the *base* scenario had the highest median number of vaccinated animals at 305,105, followed by *base x 2* with 210,786 and *base x 3* with 202,471. Interestingly, despite the higher vaccination rates, the final average number of vaccinated animals was lower in the scenarios with increased vaccination rates. This occurs because increasing the vaccination rate reduces the duration of outbreaks, ultimately resulting in fewer animals to be dosed in the latter part of the control actions period. Our findings are consistent with the results of studies conducted in Australia, Canada, New Zealand, and the United States, where emergency vaccination correlated with a reduction in the number of infected farms and a decrease in outbreak duration(54,55,55–57).

By examining the cumulative number of depopulated animals, our results showed that the *depopulation* scenario was more effective in controlling epidemics than the *base* and *base x 2* scenarios. The primary reason for its greater effectiveness was the high intensity of farm depopulation per day. Our analysis of the daily average number of depopulated animals over time revealed that the *base x 2* scenario had the highest mean cumulative count, with 3,071 animals culled, followed closely by the *depopulation* scenario, in which 2,159 animals were culled. The *base* scenario had a mean of 2,139 culled animals, whereas the *base x 3* scenario had 1,151 culled animals. When depopulation of individual species was examined, the *depopulation* scenario consistently recorded the highest number of culled animals, particularly for bovines and small ruminants. In contrast, in the *base* and *base x 2* scenarios, there was a greater prevalence of culled swine than of other species. The main reason for this is that prolonged outbreaks tend to affect more pig farms. Although there are fewer pig farms than cattle farms, the number of animals per pig farm is significantly greater (Supplementary Figure S1). While depopulation is an effective countermeasure to address highly contagious diseases such as FMD (58–60), culling healthy animals raises ethical concerns. It also increases economic losses due to reduced production and the need for farmer compensation (61). Other studies have proposed alternatives to ring depopulation; for example, (18) simulated a target density strategy and demonstrated its advantages in combating FMD; it resulted in depopulation of a smaller number of healthy animals than traditional total ring depopulation, and the time required to eliminate the outbreaks was similar. In addition, we emphasize the importance of timing when initiating depopulation for FMD control to prevent disease outbreaks across all farms in the area and potential outward spread (62). Moreover, prolonged delays in culling can lead to recurrent outbreaks in previously controlled areas, and under specific atmospheric conditions, there is a risk of long-distance airborne spread (60).

### 4.1 Limitations and further remarks

Since there have been no recent FMD outbreaks in the study region, we extracted parameters from the data on the FMD outbreak in Rio Grande do Sul (2000 and 2021) and utilized the literature to obtain the remaining parameters. Our sensitivity analysis did not identify a specific number of infected animals. Thus, our model has an acceptable level of robustness. The virulence, infectivity, and transmission of FMD can vary among strains (1,13). Although the most recent outbreaks in the State of Rio Grande do Sul involved serotypes O and A(24), we cannot rule out the possibility that other strains that exhibit different dissemination patterns might be introduced. Future work could include transmission scenarios with strains circulating in neighboring countries.

Additionally, other important between-farm transmission routes, such as vehicles and the movement of farm staff (both of which have been previously associated with FMD dissemination (63,64), were not included in our model. If such indirect contact networks are considered, the results would likely change, and the realism of the model would be improved (55). We assumed 100% compliance with the restriction on between-farm movement from infected farms and farms directly linked to infected farms and with the restriction on movement into and from control zones; we also assumed that the method by which the depopulated animals were disposed of eliminated any possibility of further dissemination of the virus. Nevertheless, real-world compliance with the control actions was not examined or considered. Our model can provide a distribution of expected FMD epidemics for any current or future control actions listed in the Brazilian control and elimination plan (65). Nevertheless, because our results are based on population data and between-farm movement data from Rio Grande do Sul, the conclusions drawn from our findings should not be extrapolated to other regions. However, according to the MHASpread model, infrastructure is highly flexible and can be easily extended to other Brazilian states and other countries.

## 5 Conclusion

In summary, we demonstrated the importance of including species-specific propagation dynamics in FMD transmission models designed to assist decision-makers in planning control and mitigation strategies for FMD. We have shown that a quick response in initiating control actions on a lower number of infected farms is crucial to reduce the necessary duration of control actions. We found that increasing depopulation capacity was sufficient to eliminate outbreaks without vaccination. Eliminating infected or likely-infected animals is an optimal strategy for preventing further epidemics, but culling large numbers of healthy animals raises welfare concerns. Regardless of which species in which FMD was introduced, the median distance over which the disease spread was within 25 km, a finding that could explain the effectiveness of the simulated countermeasures within the control areas used for FMD response. Our model projections, along with the necessary software, are available to local animal health officials. Thus, our model can be used as a policy tool for future responses to FMD epidemics through computer-based preparedness drills and capacity building and during emergency responses to FMD epidemics by providing rules of thumb generated from simulated control scenarios.

## 7 Author contributions

NCC and GM conceived the study. FPNL coordinated the data collection. NCC designed and developed computer model code, cleaned and processed the population and movement data, and performed computational analysis. FAM and CT translated scripts from R to Python with assistance from VM and AM. GM secured the funding. NCC and GM wrote and edited the manuscript. All the authors discussed the results and critically reviewed the manuscript.

## 9 Acknowledgments

Fundo de desenvolvimento e defesa sanitária animal (FUNDESA-RS).

