## Supplementary material for "Modeling foot-and-mouth disease dissemination in Brazil and evaluating the effectiveness of control measures": SS

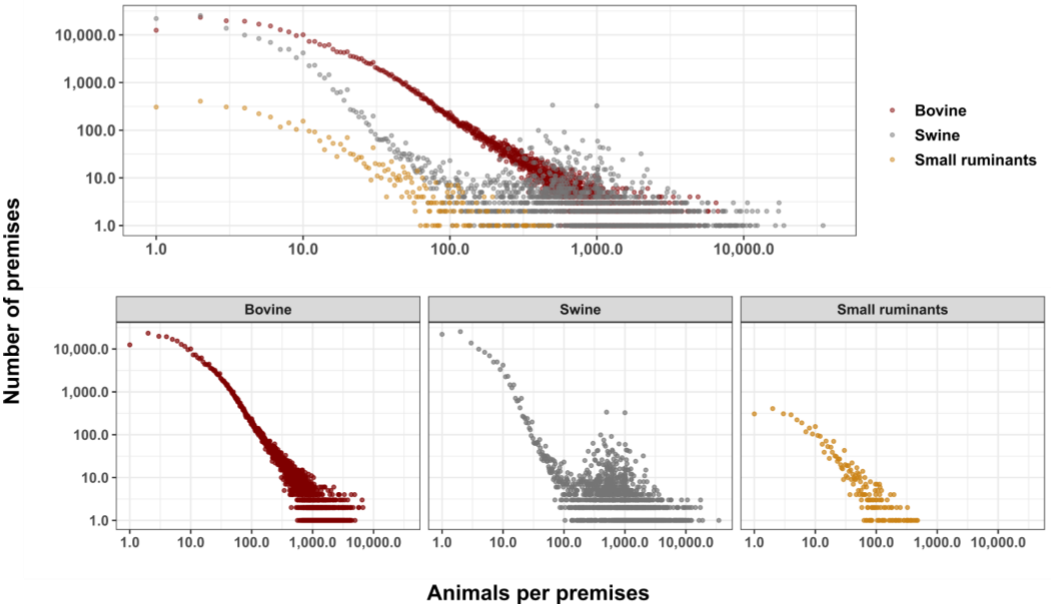


**Supplementary Figure S1.** Distribution of the farm-level populations of each species in the state of Rio Grande do Sul, Brazil. The y-axis represents the number of farms, and the x-axis represents the number of animals; both are on a log10 scale.

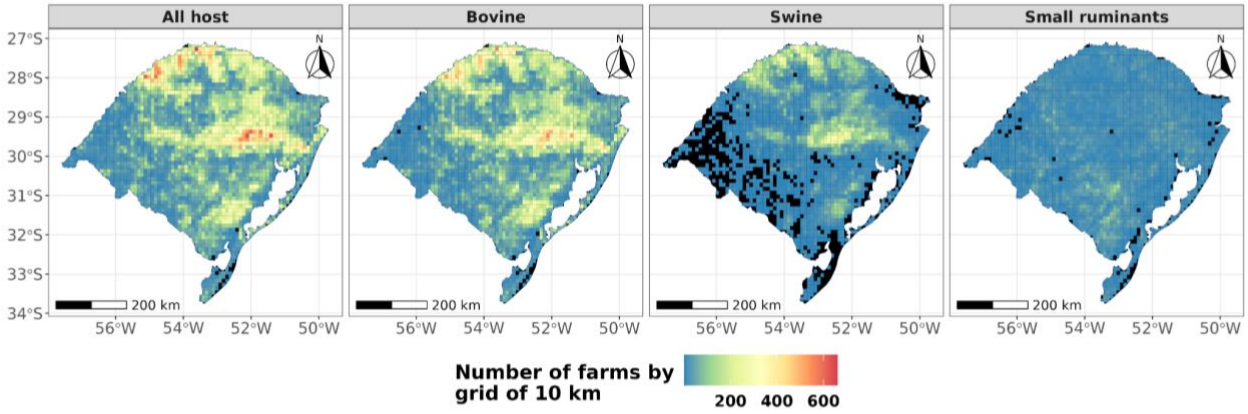


**Supplementary Figure S2.** Spatial distribution of farms in Rio Grande do Sul, Brazil. The map shows the spatial distribution of the sampled farms used to simulate the initial outbreak in the SEIR model according to the individual host species in Rio Grande do Sul over the study period.

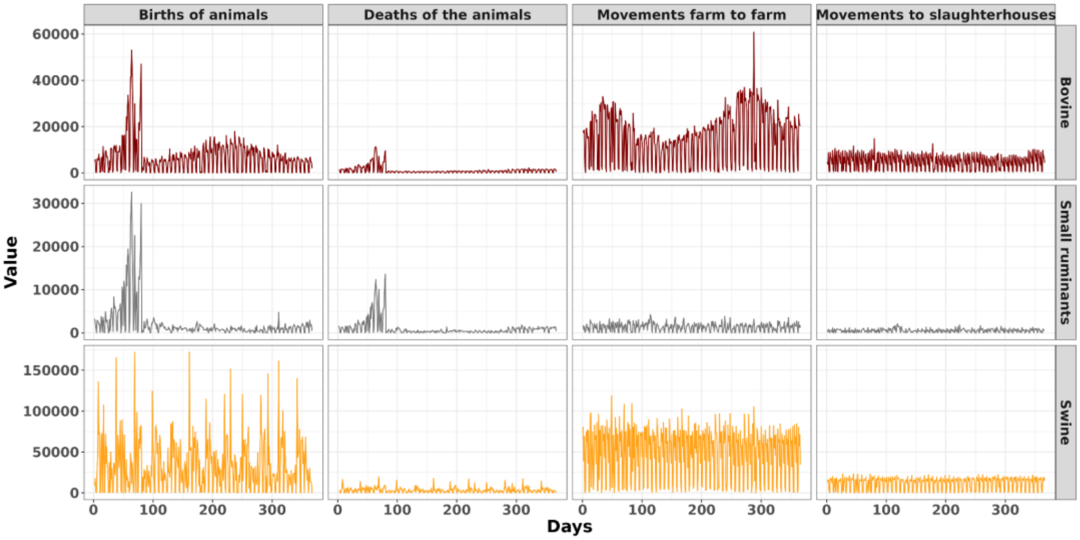


**Supplementary Figure S3.** Daily dynamics of animal movement across various transitions, including interfarm transfers, farm-to-slaughterhouse movement, births, and deaths, throughout the study period.

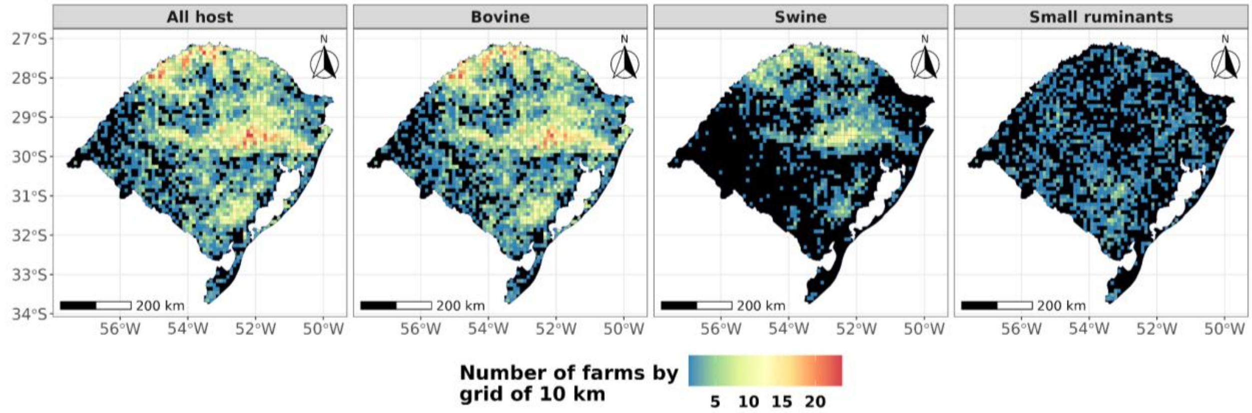


**Supplementary Figure S4.** Spatial distribution of the sampled farms used to simulate the initial outbreak in the SEIR model according to the host species in Rio Grande do Sul over the study period.

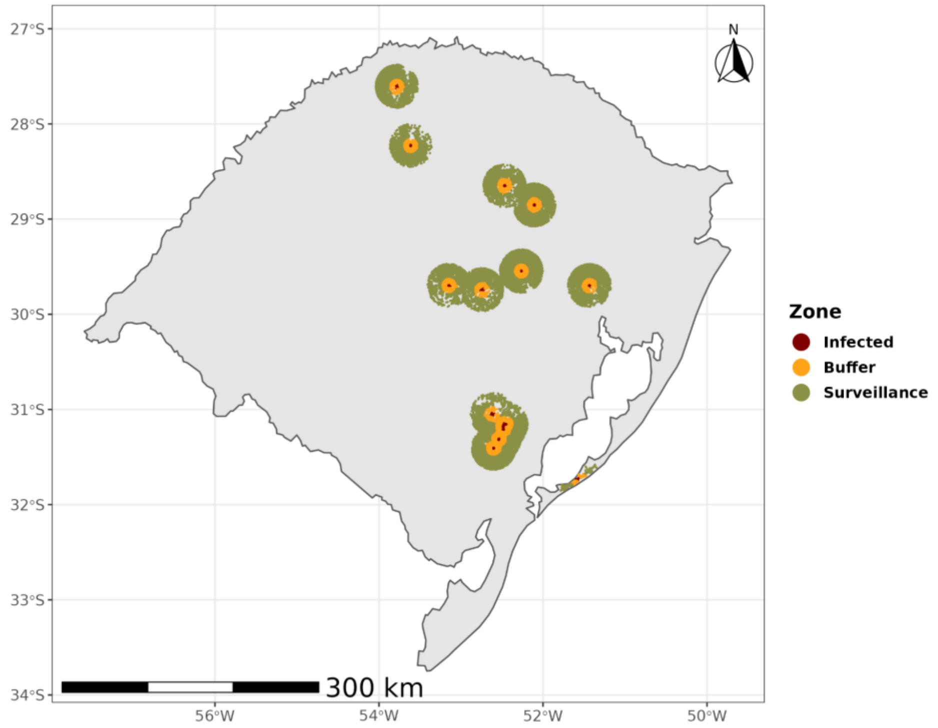


**Supplementary Figure S5**. **Control zone mapping.** The infected control zones at 3 km are represented as red dots, the 7-km buffer zone is represented as yellow dots, and the 15-km surveillance zone is represented as green dots. Notably, control zones are merged with overlapping zones.

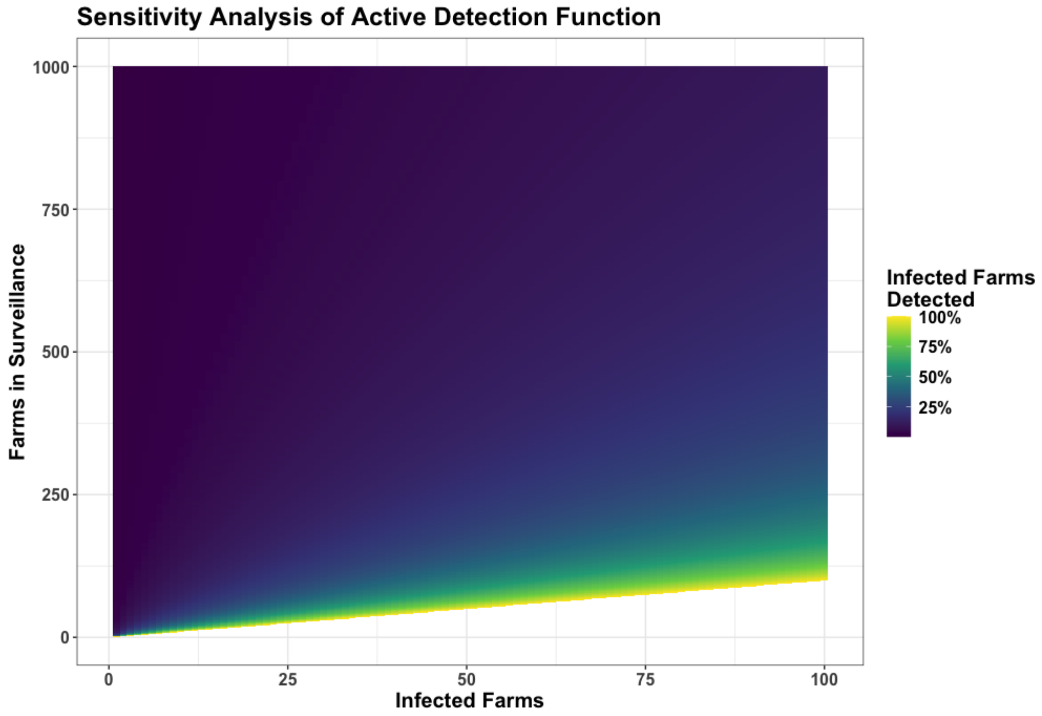


**Supplementary Figure S6. Percentages of infected farms detected according to prevalence and population size**. The y-axis represents the number of farms under surveillance, and the x-axis represents the number of infected farms in the population. The color represents the percentage of infected farms that are detected.

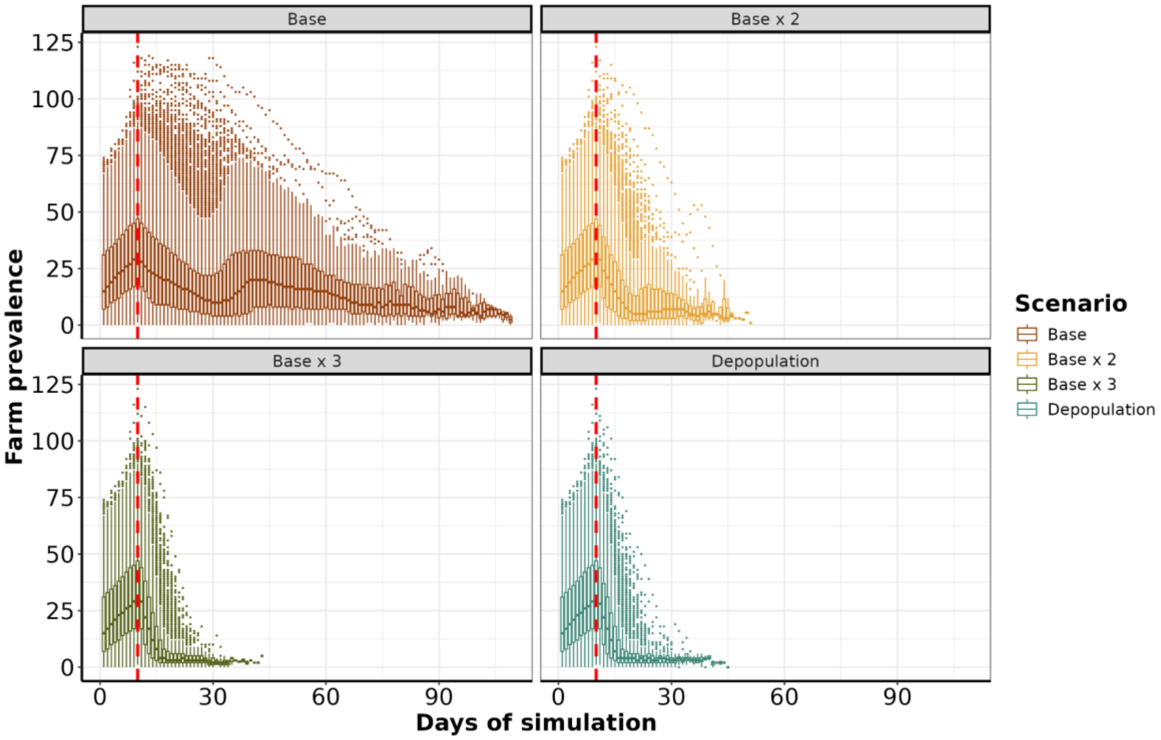


**Supplementary Figure S7. Box plot of epidemic trajectories**. The y-axis denotes the number of infected farms, and the x-axis represents the number of days in the simulation. The colors of the lines denote the control action scenarios implemented during the simulation.

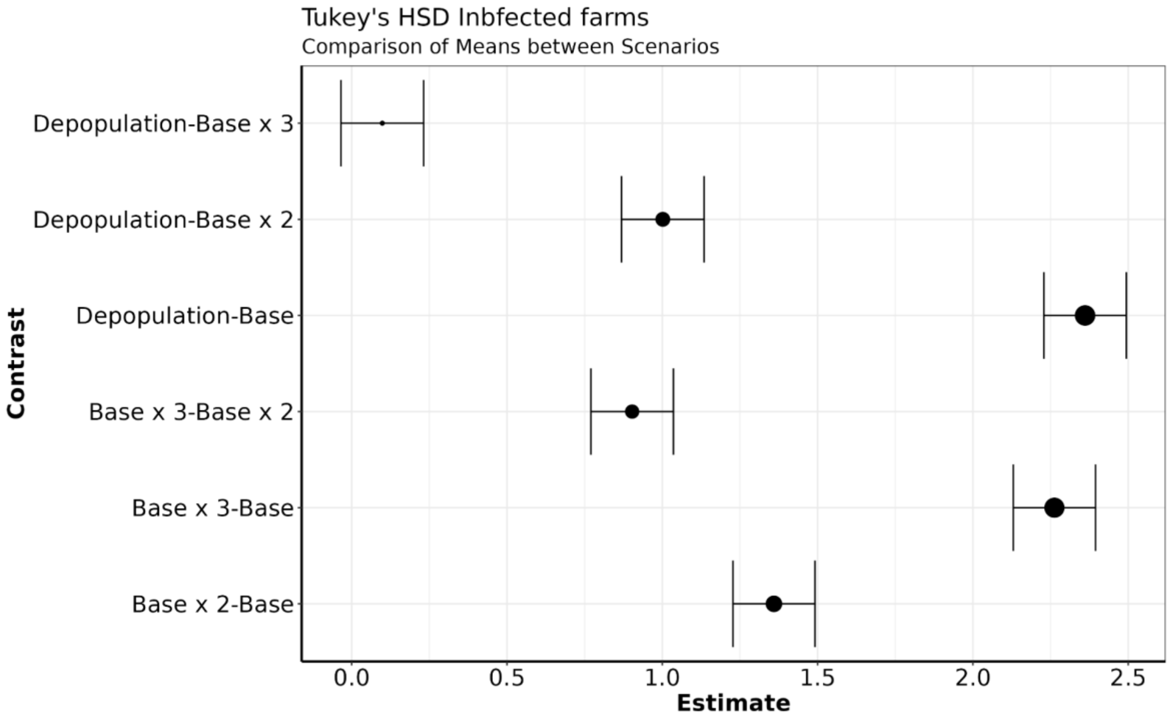

**Supplementary Figure S8. Tukey multiple comparisons of the mean numbers of infected farms**. Comparison between scenarios at 95% familywise confidence levels across different scenarios are shown.

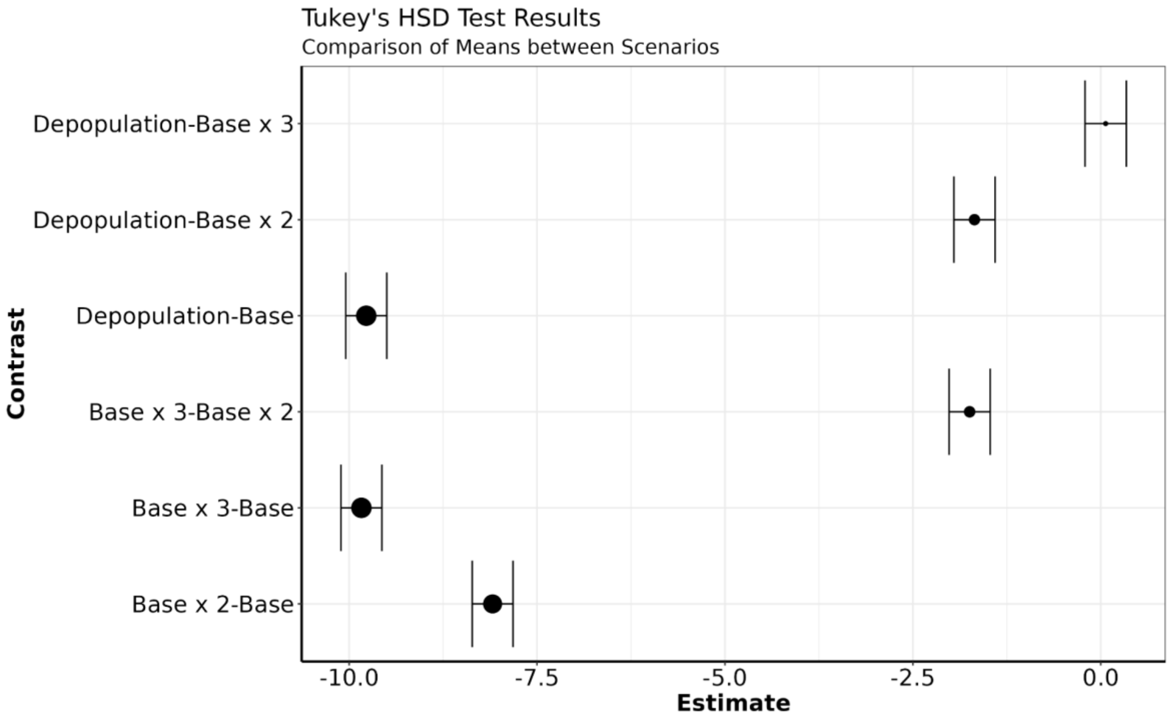


**Supplementary Figure S9. Tukey multiple comparisons of the mean numbers of days spent working on control actions**. Comparisons between scenarios at 95% familywise confidence levels are shown.

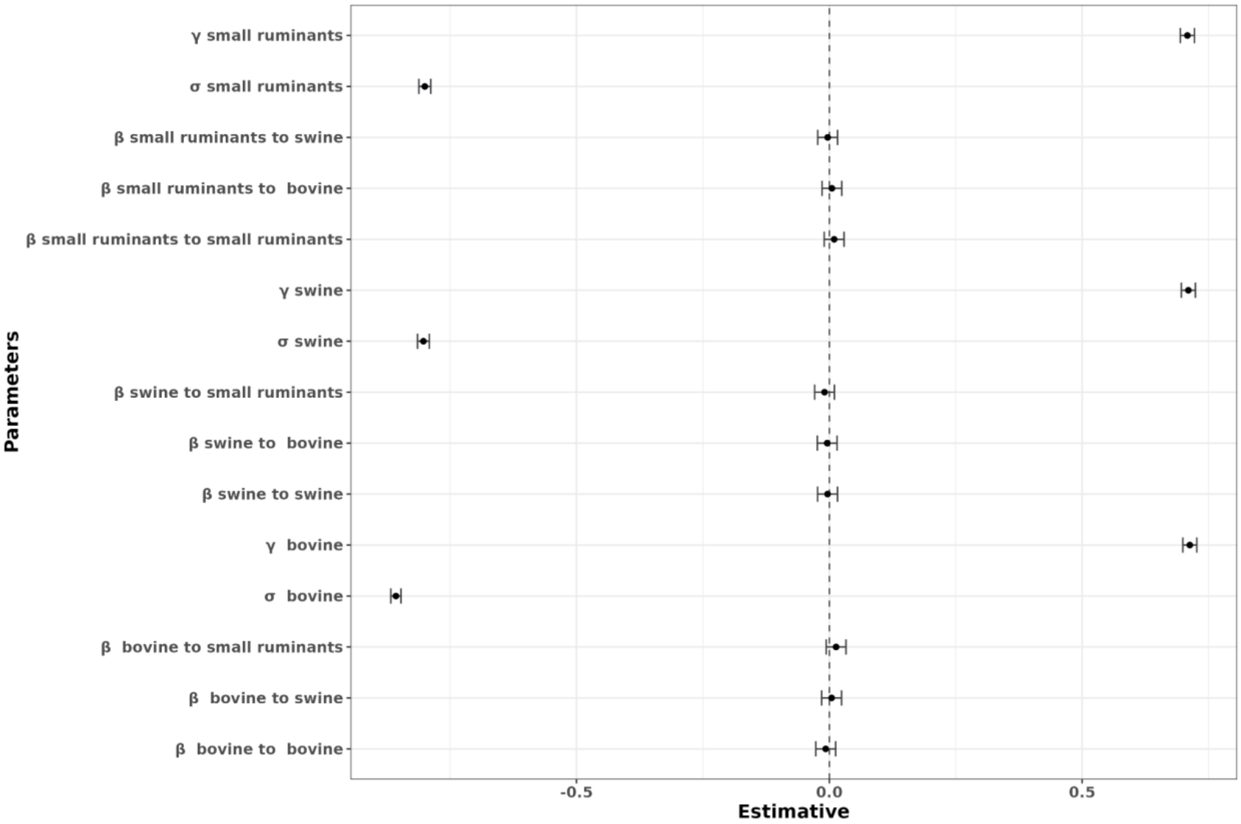


**Supplementary Figure S10.** Sensitivity of the number of infectious animals to changes in the parameters via the LHS-PRCC.

### 1 Supplementary Material Methods

**Mathematical description of the initial spread model**

Here, we described one interaction between bovines and swine; however, the equations can be easily adapted to describe SEIR dynamics involving other species. We denote the compartments for each species within a specific farm as $S_i, E_i, I_i$, and $R_i$ and define subscripts $i$ and $j$ such that $i, j = 1$ for bovine, $i, j = 2$ for swine, and $i, j = 3$ for small ruminants.

The equations describing the bovine populations of individual farms can be written in the following manner:

1. Susceptible population $(S_1)$:

$$\frac{dS_1}{dt} = \Lambda_1 S_1 - \beta_{12}\frac{S_1 \cdot I_2}{N_1} - \beta_{13}\frac{S_1 \cdot I_3}{N_1} - \mu_1 S_1 + \gamma_1 R_1$$

2. Exposed population $(E_1)$:

$$\frac{dE_1}{dt} = \beta_{12}\frac{S_1 \cdot I_2}{N_1} + \beta_{13}\frac{S_1 \cdot I_3}{N_1} - \sigma_1 E_1 - \mu_1 E_1$$

3. Infectious population $(I_1)$:

$$\frac{dI_1}{dt} = \sigma_1 E_1 - \gamma_1 I_1 - \mu_1 I_1$$

4. Recovered population $(R_1)$:

$$\frac{dR_1}{dt} = \gamma_1 I_1 - \mu_1 R_1$$

where $\Lambda_i$ = the birth rate of the animals of species $i$ within that particular farm, $\mu_i$ = the death rate of the animals of species $i$ within that particular farm, $\beta_{ij}$ = the transmission rate from species $j$ to species $i$, $\sigma_i$ = the rate of transition of animals from exposed to infectious for species $i$, and $\gamma_i$ = the recovery rate for species $i$.

$N = S + E + I + R$ = Total population size. $t$ = represents time and is defined on a daily timescale.

**Control action modeling**

1. *Vaccination strategy modeling*

The vaccination procedure described below is implemented within the control action modeling dynamics in a population of farms. *Here*, the simulation model operates on population data for the animal population within each farm.

*Vaccination strategy*

The vaccination strategy involves targeting a specified number of farms in the selected area for vaccination each day. The vaccination strategy takes into account the following:

- The number of farms to be vaccinated per day is determined on the basis of the proposed scenario. In this scenario, vaccination will be administered to farms located within the infected zone and/or the buffer zone according to predefined criteria. The priority of farms vaccination will be sorted from highest to lowest on the basis of the number of animals in the population.

- The day on which vaccination at a specific farm is initiated; after 15 days, vaccination is no administered at that farm.
- The number of days required for 90% immunity to develop across the vaccinated farm population.

We selected the animals to be vaccinated as follows:

$$n = (1 - ve) * N$$

$ve$ = 0.9, meaning that the vaccine has an efficacy of 90%; in other words, it is expected to be effective in preventing disease in 90% of the vaccinated animals. We select how many of this number of animals will be moved to vaccinated status via a binomial distribution with a probability of $vt$.

Thus,

$$X \sim Binomial(n, p)$$

where $p = 1/vt$ and $vt$ represents the 15 days needed to assume that a given animal will be fully immunized.

#### 2 Infected farms detection algorithm

The process used in the algorithm is similar to a method for simulating surveillance efforts on farms, taking into account the number of infected farms and the total number of farms under surveillance.

Let us denote:

-$P_i$: Number of farms under surveillance.
- $I_i$: Number of infected farms.
- $E$: Number of farms found in the current iteration

The algorithm operates as follows:

1. If $I_i < 5$, $E$ is set to a random value between 0 and $I_i$ (inclusive), i.e., $E = sample(0{:}\, I_i, 1)$.

2. If $I_i \geq 5$ a probability distribution is calculated via the hypergeometric distribution with parameters ( $m = I_i$), ( $n = P_i$) and ( $k = \frac{p_i}{3}$ ).
Sample a value $p$ from this probability distribution.
Calculate $E = [I_i * p]$
If $E = 0$, set $E = 1$

To illustrate the algorithm's behavior, let us consider an example in which we simulate the portability of infection across a spectrum of scenarios. We examined the impact of the number of infected farms over a range of from one to one hundred and the impact of populations ranging in size from ten animals to one thousand animals. The results are depicted in Supplementary Material Figure S6.
